# High-throughput enrichment and isolation of megakaryocyte progenitor cells from the mouse bone marrow

**DOI:** 10.1101/512442

**Authors:** Lucas M. Bush, Connor P. Healy, James E. Marvin, Tara L. Deans

**Affiliations:** Department of Biomedical Engineering, University of Utah, Salt Lake City, UT 84112, USA; Flow Cytometry Core Facility, University of Utah Health Sciences Center, Salt Lake City, UT 84112, USA

**Keywords:** Megakaryocytes, hematopoietic stem cells, image flow cytometry, myeloid bypass, principal components analysis

## Abstract

Megakaryocytes are a rare population of cells that develop in the bone marrow and function to produce platelets that circulate throughout the body and form clots to stop or prevent bleeding. A major challenge in studying megakaryocyte development, and the diseases that arise from their dysfunction, is the identification, classification, and enrichment of megakaryocyte progenitor cells that are produced during hematopoiesis. Here, we present a high throughput strategy for identifying and isolating megakaryocytes and their progenitor cells from a heterogeneous population of bone marrow samples. Specifically, we couple thrombopoietin (TPO) induction, image flow cytometry, and principle components analysis (PCA) to identify and enrich for megakaryocyte progenitor cells that are capable of self-renewal and directly differentiating into mature megakaryocytes. This enrichment strategy distinguishes megakaryocyte progenitors from other lineage-committed cells in a high throughput manner. Furthermore, by using image flow cytometry with PCA, we have identified a combination of markers and characteristics that can be used to isolate megakaryocyte progenitor cells using standard flow cytometry methods. Altogether, these techniques enable the high throughput enrichment and isolation of cells in the megakaryocyte lineage and have the potential to enable rapid disease identification and diagnoses ahead of severe disease progression.

## Introduction

Megakaryocytes (MKs) are important cells within the hematopoietic lineage because they produce platelets, the anucleate cells that facilitate healing. Platelets are released into the bloodstream from MKs where they play a central role in homeostasis, immune responses, and hemostasis^1,2^. Platelets also play an important role in thrombosis, which underlies heart attacks and stroke, making them the target of many pharmaceutics and therapies. MKs differentiate from hematopoietic stem cells (HSCs)^3^, but the mechanisms of MK and subsequent platelet development are not fully understood, leading to difficulties when identifying the circumstances that cause disease^4–7^. MK maturation is characterized by an increase in cell size and DNA content to develop multilobed polyploid nuclei, an upregulation of MK-specific surface markers, and cytoplasmic remodeling to prepare for platelet production^5,8^. A significant challenge with studying MK development, and the diseases that arise from disrupted MK development, is the identification and classification of the progenitor cells that are produced from HSCs and subsequently give rise to MKs^9^. Although MKs are continually produced through adulthood, HSCs and MKs account for only 0.5% and 0.01% of cells, respectively, in the mouse bone marrow. The frequency of MK progenitors is expected to be similarly infrequent which presents significant challenges for large-scale studies of this intermediate cell population^10,11^.

Recent studies suggest that MKs are derived from HSCs through multiple parallel lineage pathways^12–15^. While these lineage pathways are not fully elucidated, their diversity highlights the need to identify and isolate MK progenitor cells in order to understand MK development and maturation. However, because the bone marrow contains a heterogenous population of cells, it is difficult to identify MK progenitor cells. The standard approaches to distinguish individual cells in the hematopoietic lineage is to use combinations of antibodies that recognize cell-specific surface markers and identify cells based upon their immunofluorescent profiles when analyzed by flow cytometry. In the mouse, HSCs are broadly defined by the Lin^−^ Sca-1^+^ c-Kit^+^ (LSK) immunophenotype^16,17^, and have two distinct populations: 1) HSCs that are capable of long-term bone marrow reconstitution (LT-HSCs) and 2) HSCs capable of short-term hematopoietic reconstitution (ST-HSCs). Two different differentiation pathways, classical and alternative, have been proposed where ST-HSCs give rise to MK cells in which precursor cells progress through multiple differentiation states in a step-wise fashion before becoming a MK cell^17^ (Fig. 1A,B). Notably, in neither of these pathways are the MK progenitor cells thought to have any proliferative potential. In contrast, the recently proposed myeloid bypass model suggests that a population of self-renewing MK repopulating progenitor (MKRP) cells are formed directly from LT-HSCs (Fig. 1C). The myeloid bypass model suggests that the formation and development of MKs may be dynamic and amenable to ever-changing biological needs^15^. Identifying and isolating this MKRP cell population holds the potential to rapidly advance our understanding of MK development and maturation.

**Figure 1:**
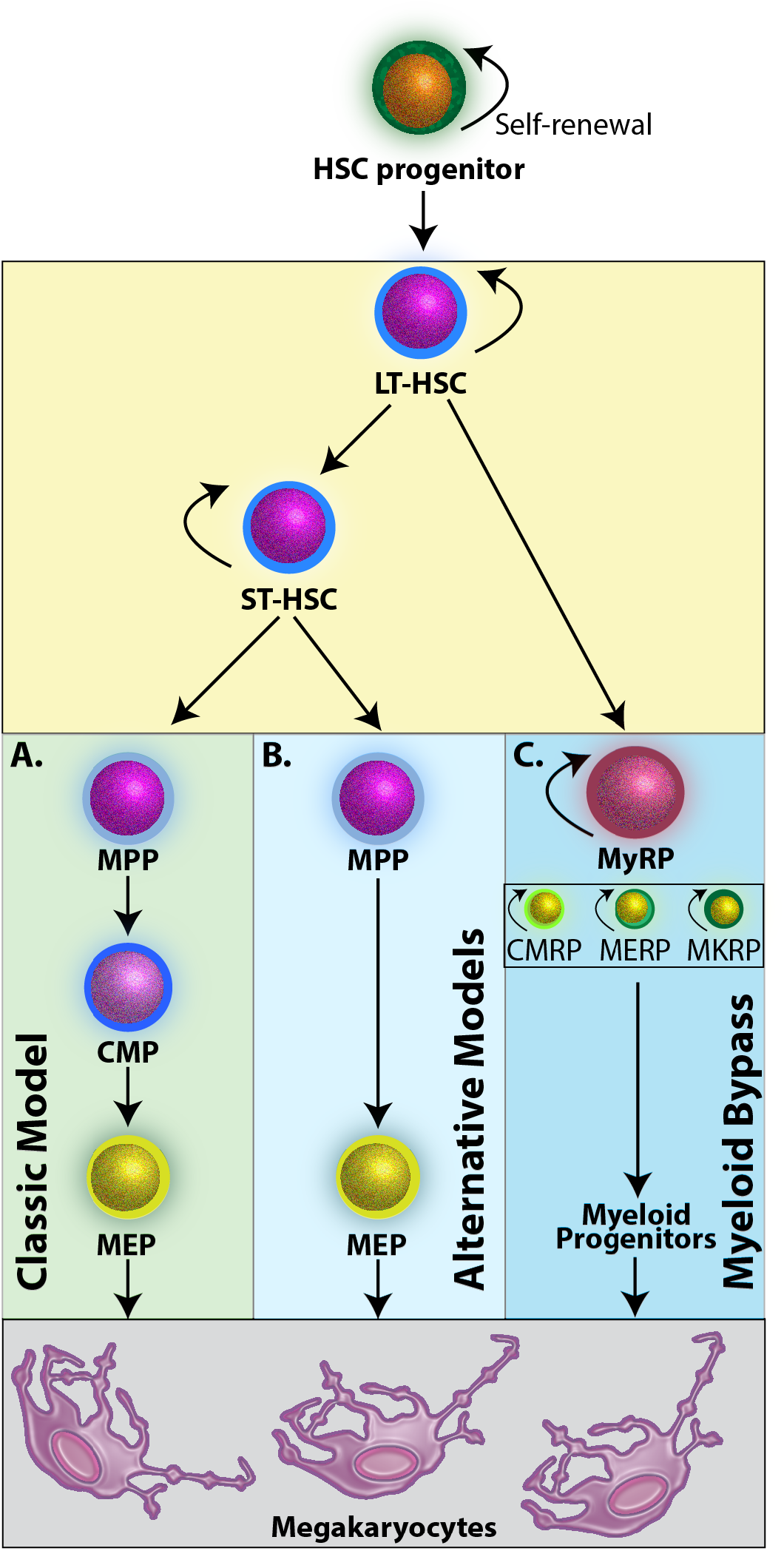
Models of MK differentiation from HSCs. **A.** In the classic model, cell-fate decisions progress in a step-wise fashion where ST-HSCs progress toward an MPP cell population that has no self-renewal capacity but can further commit to CMP cells, MEP cells, and finally MKs. **B.** The alternative model yields MEP cells directly from non-self-renewing MPPs. **C.** The Myeloid bypass model suggests that LT-HSCs are primed to directly give rise to myeloid progenitor cells that all have the capacity to self-renew and maintain progenitor populations. Abbreviations: HSC, hematopoietic stem cell; ST-HSC, LT-HSC, long-term hematopoietic stem cell; short-term hematopoietic stem cell; ST-HSC, short-term hematopoietic stem cell; MPP, multipotent progenitor; CMP, common myeloid progenitor; MEP, megakaryocyte/erythroid progenitor; MK, megakaryocyte; MyRP, myeloid repopulating progenitor; CMRP, common myeloid repopulating progenitor; MERP, megakaryocyte-erythroid repopulating progenitor; MKRP, megakaryocyte repopulating progenitor.

Because most cells develop distinct morphological characteristics at various stages of differentiation, we hypothesize that cellular morphology may be an important parameter to identify the MKRP cell population. Consistent with this hypothesis, commitment down the MK lineage is characterized by an increase in cell size, endoreplication that increases DNA content, and cytoplasmic remodeling to prepare for platelet production^5,8^. These individual morphological changes, along with fluorescently-labeled cell markers, can be observed using image flow cytometry, where an image of each individual cell is captured as it flows through the cytometer^18^. Each captured image contains data of multiple variables that can be used to characterize individual cells by cell size, shape, and membrane texture, as well as expression levels of fluorescently-labeled cell surface markers. These characteristics can be quantified using computational approaches like principal components analysis (PCA). PCA is a statistical method that groups highly correlated data, such as phenotypic characteristics, by converting a set of possibly correlated variables in a multivariate dataset into sets of values called principal components. Principal components allow multidimensional observations to be grouped based upon overall similarity in order to identify trends in datasets that would otherwise be intractable due to their high dimensionality. In short, this method captures variance in a multivariant dataset to identify the most correlated features that can be used to distinguish cells from different stages of lineage development. Here, we couple conventional antibody labeling, image flow cytometry, and PCA analysis to identify a repopulating population of MK progenitor cells. By coupling these analyses with a cell culture method that enriches for MKRP cells we can reliably identify these cells from heterogenous bone marrow samples using both image flow and conventional flow cytometry. We envision that this will facilitate studies leading to a better understanding the development of MKRP cells, in addition to approaches for increasing MK and platelet production for disease treatment.

## Results

### Image flow cytometry analysis of bone marrow cells following TPO-mediated expansion

MKRP cells often express canonical MK markers that are similar to other myeloid cells, making it difficult to distinguish early progenitor subpopulations from other hematopoietic cells without using large antibody panels. Since most cells develop distinct morphological differences when they are at various stages of cell fate, cell morphology is an additional parameter that can be used to identify MKRP cells. Image flow cytometry is an ideal tool for identifying cell populations and distinct stages of lineage development based upon these differences because it enables the simultaneous analysis of morphologic characteristics and cell surface marker expression.

To identify the cells that participate in the commitment to MK cells, mouse bone marrow cells were harvested and characterized by the expression of c-Kit, Sca-1, and lineage (Lin) surface markers. This immunofluorescence labeling strategy, known as LSK staining (Lin^−^ Sca-1^+^ c-Kit^+^), is historically used to identify HSCs that can be distinguished by analysis using flow cytometry (Fig. 2A, Supplementary Methods). The TPO/c-Mpl signaling within the hematopoietic niche of the mouse bone marrow is essential for HSC survival, proliferation and differentiation into mature MKs^5,8,12,19–22^. Therefore, the TPO cytokine is commonly used for stimulating and differentiating HSCs to become MKs^23^. The cells harvested from the bone marrow and grown in TPO for 72 hours experience a shift in Lin, Sca-1, and c-Kit expression (Fig. 2C, Supplementary Fig. 1–3), indicating these cells are primed for differentiation. Comparing image files associated with cells grown in each growth condition shows drastic morphological changes of the cells in the R4 subpopulation of cells (Fig. 2B,D). Cells in the R4 subpopulation are substantially larger and have an increased in Sca-1 and c-Kit expression after exposure to TPO. These cells are present in both growth conditions, but enrichment upon TPO treatment suggests that TPO may prime the cells in the R4 subpopulation for lineage commitment into MKs.

**Figure 2:**
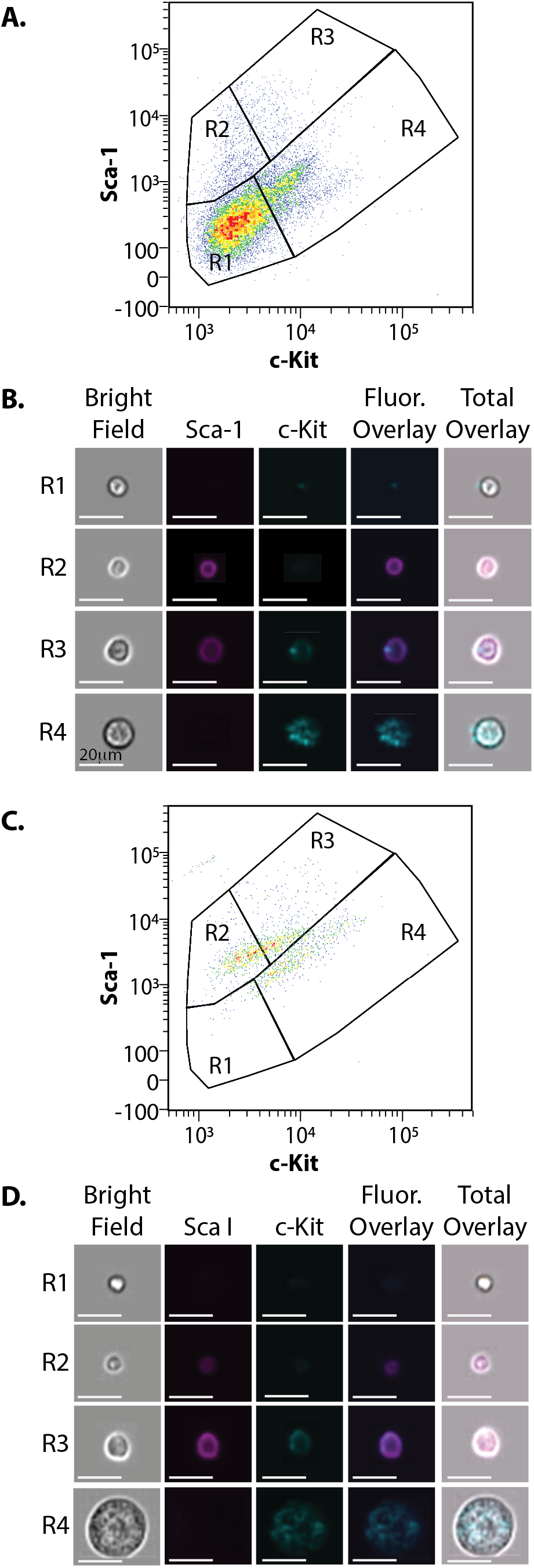
Characterization of progenitor cell subpopulations of mouse bone marrow cells. **A.** Cells were harvested from the mouse bone marrow, stained with LSK antibodies and run on an image flow cytometer. Cells were gated for Lin^−^ cells and the total population was divided into four distinct subpopulations (R1-R4). **B.** As cells ran through the image flow cytometer, an image was captured of each cell. Individual cells of each of the four subpopulations are represented. **C.** Bone marrow cells grown in the presence of TPO then stained with LSK antibodies were run on the image flow cytometer. Cells were divided into four distinct subpopulations (R1-4). **D.** Representative image panels of each subpopulation after cells are grown in culture with 50 ng/mL of TPO and stained with LSK antibodies. Scale bars for all images, 20μm.

### Principle component analysis identifies MKRP cells

Because HSCs have been shown to differentiate into MK cells in the presence of TPO, we aimed to use PCA to identify transitions in cell morphologies to locate the MKRP cells. To validate that distinct subpopulations of cells can be characterized morphologically, PCA was used to mathematically capture the variance in image datasets obtained from the image flow cytometer for both the −TPO and +TPO growth conditions stained with the LSK immunofluorescence panel. With PCA, the input variables contribute to each principal component such that the first principal component explains the largest possible variance in the dataset^24^. The closer a given cell’s principal component score is to another cell’s principal component score, the more similar those two cells are morphologically. This approach determined the most correlated morphological characteristics of each cell using the bright field images for cells directly isolated from the bone marrow that were not treated with TPO (Fig. 2B) and cells cultured with TPO for 72 hours (Fig. 2D). Twenty-nine morphological features corresponding to the shape, size, and texture of the cells in the dataset were considered in the principal component analysis (Supplementary Fig. 4A-C). Analyzing the percent variance of each principal component with the broken stick model, which determines two principal components are needed to effectively interpret the data, shows that in the −TPO dataset, PC1 (size) explained 60% of the morphological variation between cells while PC2 (shape) explained 14% of the morphological variation between cells, both exceeded the broken stick model (Fig. 3A, Supplementary Methods)^25^. In the +TPO dataset, only PC1 (size) exceeded the broken stick model but PC1 also explained over 75% of the morphological variation in the dataset (Fig. 3B). Therefore, TPO treatment enhanced the morphological similarity of cells. In both datasets, PC1 correlated most highly with the size parameters: diameter and area (Fig. 3C,D). This suggests that the morphology of cells in both datasets varied most widely based on these two categories while less variation was observed in other morphological features.

**Figure 3:**
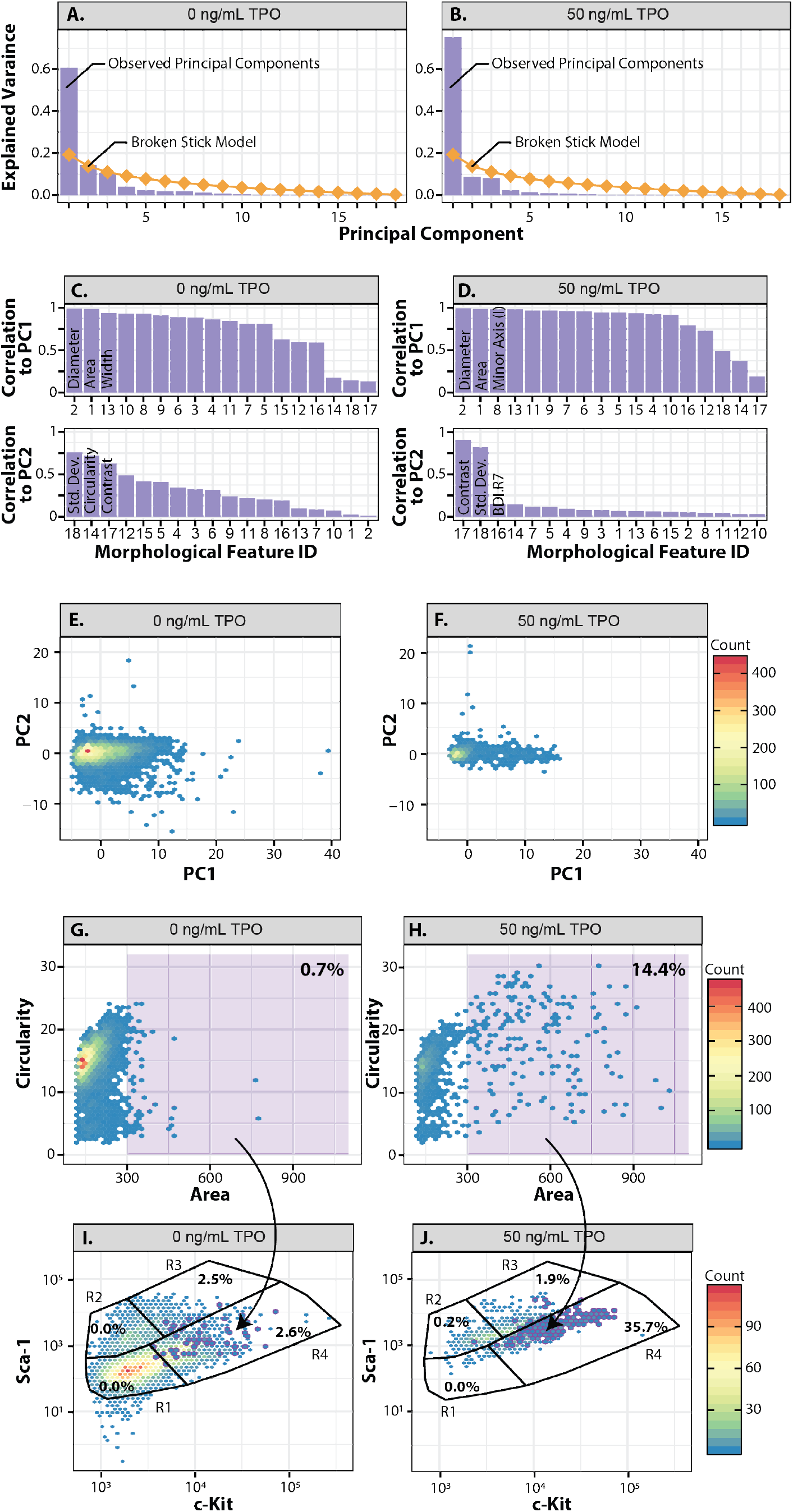
Morphology predicts MKRP phenotype using PCA. **A.** Scree plot showing percent eigenvalue for each principal component in the LSK morphology dataset with and **B.** without TPO. The theoretical percent variation under the broken-stick model is represented as a diamond dashed line, which indicates the boundary between interpretable and uninterpretable principal components. **C-D.** The absolute contribution/correlation of each morphological feature to each morphological principal component **C.** without TPO and **D.** with TPO. **E-F.** The top three most highly correlated features are listed for each principal component. Color indicates the type of feature (green represents size, pink represents shape, and blue represents texture). **E.** Projections of LSK cells grown in the absence TPO and **F**. the presence of TPO on the space defined by the first and second principal components. **G.** Cell area vs. circularity of LSK cells grown in the absence of TPO and **H.** grown in the presence TPO. A gate is included (shaded purple) to mark particularly large and circular cells with the percent of cells present in each gate. **I-J.** PCA data mapped back onto the experimental flow data in the **I.** absence of TPO and **J.** presence of TPO.

Next, PC1 and PC2 from the two growth conditions were projected onto a plot onto a plot and shows clear morphological differences between the R4 subpopulation and the other subpopulations of cells as they undergo differentiation (Fig. 3D,E, Supplementary Fig. 4D,E). The cells in the R4 subpopulation are also extended along the PC1 axis and are largely distinguished by their unique size, and their morphological differences within this subpopulation of cells, which lose much of their variation in shape after growing in TPO to become a more homogeneous population of cells (Fig. 3E,F). Using PCA to analyze circularity vs. area shows that the R4 subpopulation of cells is the largest, most circular cell type, a property that is exaggerated after culturing with TPO (Fig. 3G,H). Mapping the PCA data back onto the flow data indicates that the R4 subpopulation of cells is likely a MKRP population of cells (Fig. 3I,J).

Projecting the principal component scores of each cell from the two growth conditions onto a plot shows that TPO treatment compresses the morphological variation in the cell population (Fig. 3E,F). Furthermore, the remaining variation in the TPO treated dataset is primarily due to differences in size (Fig. 3H). Overall, PCA revealed that cell size, particularly in TPO treated populations, may be sufficient to distinguish cells in the R4 subpopulation. Applying the results of PCA analysis by plotting cell area versus circularity and gating for cells with areas greater than 300, we demonstrate that TPO treatment increases the size of a specific subpopulation of cells and that this subpopulation belongs primarily to the R4 subpopulation of MKRP cells (Fig. 3G,H). Because MK maturation is marked by an increase in cell size^5^, and high levels of c-Kit expression indicate an intrinsic MK bias^26^, we hypothesized that the R4 subpopulation of cells is a LT-HSC-derived MKRP subset of cells capable of self-renewal and differentiating into mature MKs.

### Flow cytometry analysis of maturing bone marrow cells

To investigate the PCA prediction that MKRP cells exist in the R4 subpopulation, mouse bone marrow cells were harvested and analyzed before and after culture with TPO. The cells not grown in TPO were directly harvested from the bone marrow and immediately washed, then stained with antibodies to identify MKRP cells^27^. The population of cells exposed to TPO were grown in 50 ng/mL of TPO for 72 hours before being stained with the same panel of antibodies. Flow cytometry was used to assess the expression of surface markers in both growth conditions (Fig. 4A,B, Supplementary Methods). Initially, a gate was drawn based on the Lin, Sca-1, and c-Kit expression to identify the R4 subpopulation of cells identified by the PCA analyses as being the potential MKRP population of cells in both growth conditions (Fig. 4A,B). To determine if this population of cells are undergoing differentiation and are of HSC origin, a gate was drawn to include CD9 positive and Thy1.1 negative cells. To confirm that these cells were committing to the MK lineage and not the B-cell lineage, the percentage of cells that were IL7Rα negative and CD41 positive were identified (Fig. 4A,B). Comparing the percentage of MKRP cells in each condition, we showed that a subset of the R4 subpopulation likely contains MKRP cells and can be enriched when cultured with TPO (Fig. 4C,D).

**Figure 4:**
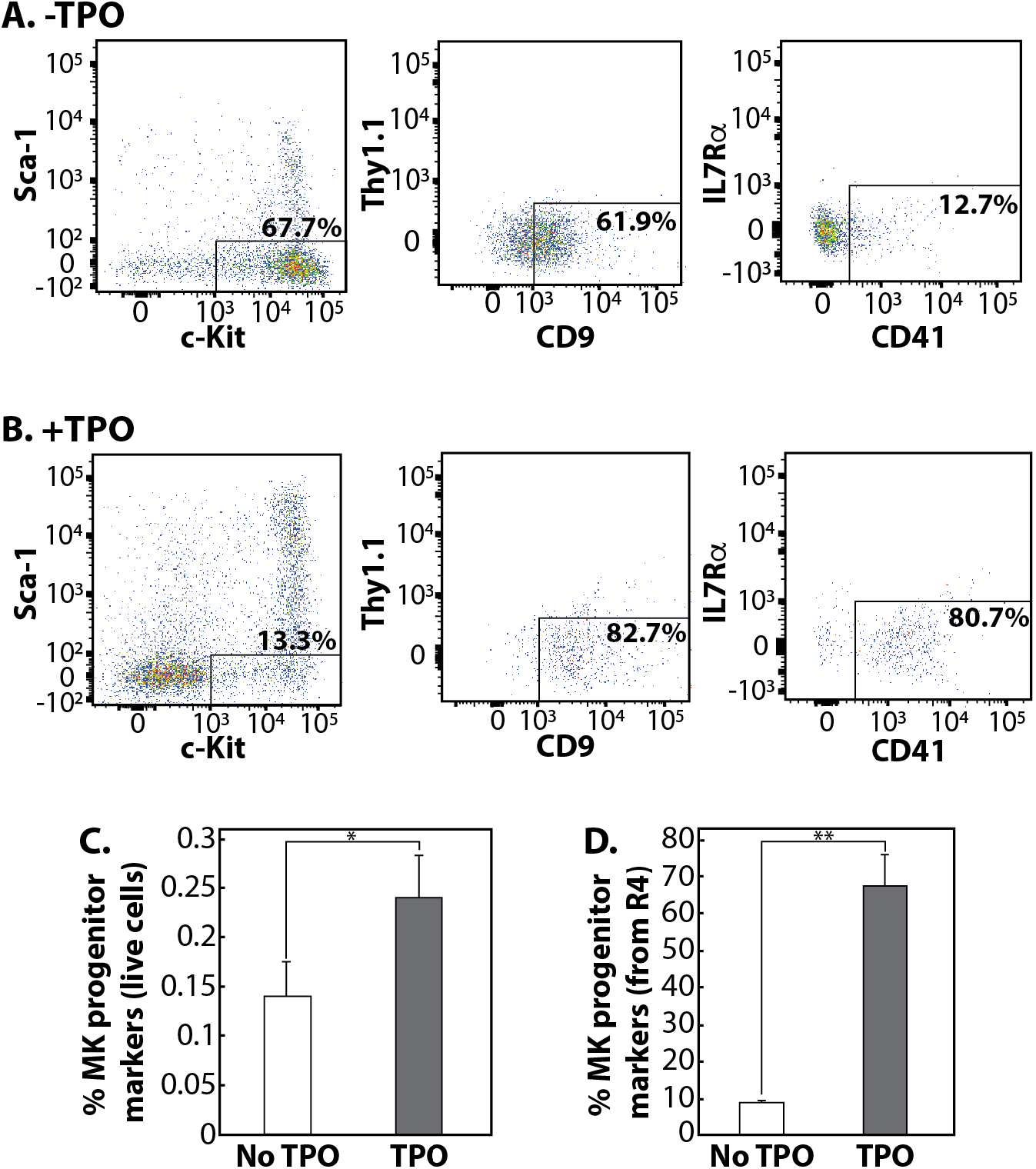
A subset of bone marrow cells are MKRP cells. **A.** Representative flow cytometry panel of fresh bone marrow cells not grown in TPO after gating on live, lineage negative cells. Boxes represent gates to identify the potential MKRP cells. **B.** Representative flow cytometry panel of harvested bone marrow cells grown in TPO after gating on live, lineage negative cells. Boxes represent gates to identify the potential MKRP cells. **C.** The percentage of R4 live cells not grown in TPO that stain for MKRP cell markers. **D.** The percentage of R4 cells that stain for MK progenitor cell markers. For **C** and **D**, the percent of specified cells represents the average from three independent mice. The error bars represent the standard deviation of the mean. *p<0.05; **p<0.005.

### MKRP cells become mature MKs

To further classify the potential MKRP cell population, we investigated the development and maturation of these cells. To observe the transition from HSCs to MKs, mouse bone marrow cells were harvested and analyzed before and after exposure to TPO. The population of cells not grown in TPO were immediately washed and stained with MK antibodies (Supplementary Table 1) after being harvested from the bone marrow, and image flow cytometry was performed to assess the expression of the MK surface markers (Fig. 5A, Supplementary Methods)^28^. The cells cultured with TPO were isolated from the bone marrow, washed, and plated in media with TPO for 72 hours before being stained with the same panel of MK antibodies (Fig. 5B). In both experimental cases, flow cytometry analysis revealed seven subpopulations of cells (R1-R7) with distinct surface marker expression levels (Fig. 5A,B, Supplementary Fig. 5). In response to TPO, the cell subpopulations shifted (Supplementary Fig. 6) and increased in MK surface marker expression levels, indicating cells differentiating from a multipotent state HSC state to a more committed MK state.

**Figure 5:**
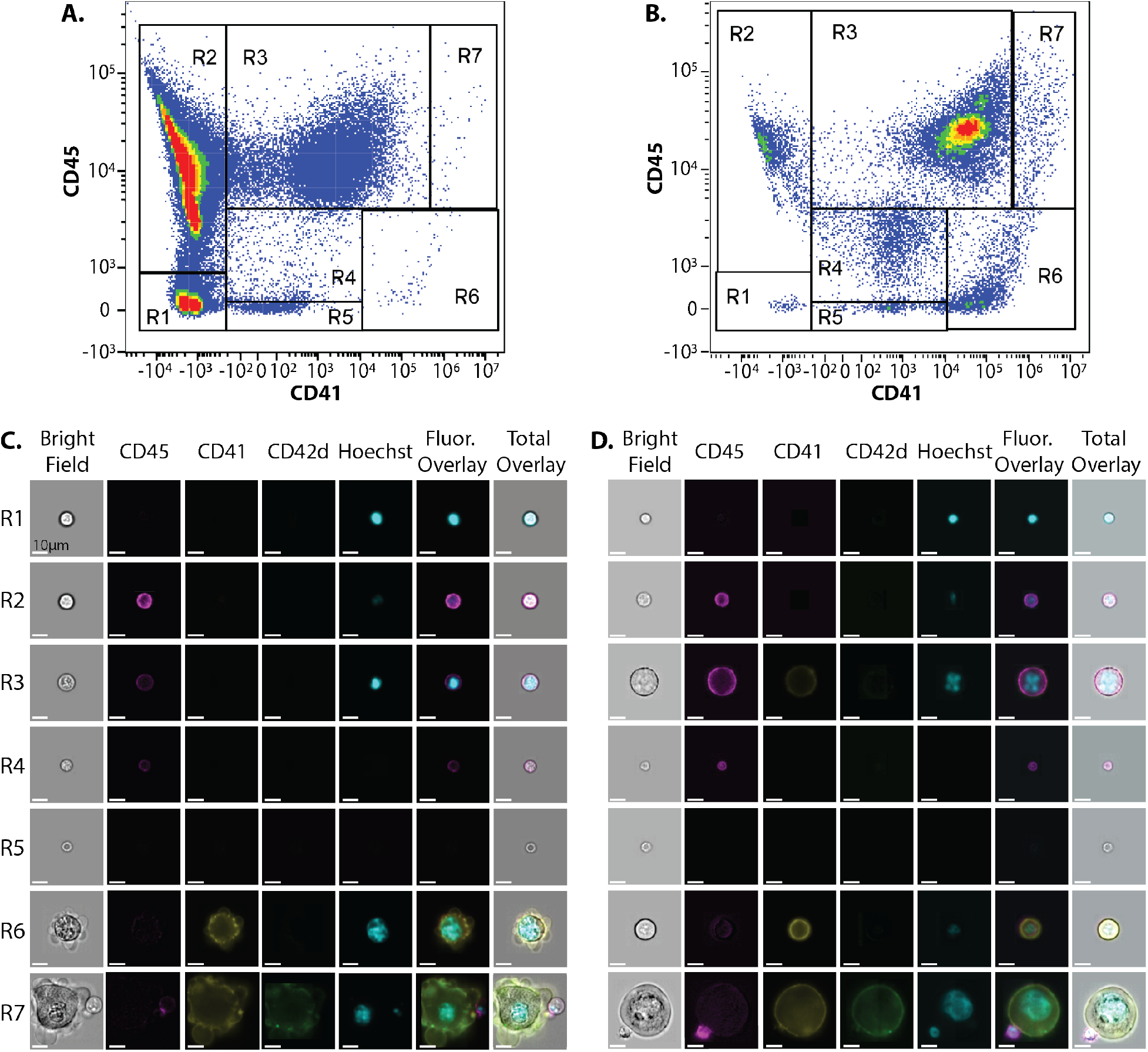
Classification of MKRP subpopulations. **A-B.** Flow cytometry plots displaying CD45 and CD41 intensity of bone marrow cells stained with MK antibodies **A.** before and **B.** after culturing with TPO. Seven distinct subpopulations were identified (R1-7). **C.** Representative images of cells before TPO and **D.** after culture in 50ng/ml of TPO. Scale bars, 10μm for all images.

Qualitative inspection of image files associated with individual flow cytometric events led to the observation that cells in the R7 subpopulation of cells were exceptionally large with the co-expression of MK surface markers and multilobed, polyploid nuclei within the confines of the cell membrane (Fig. 5C,D). To quantify this, the mean fluorescence intensity (MFI) of the three MK surface markers and the total area of Hoechst 33342 signaling were calculated for each subpopulation, revealing that the subpopulation of cells in the R7 gate expressed the highest levels of all three surface markers and had the largest nuclear area of the seven subpopulations (Supplementary Fig. 7). When plotting brightfield circularity versus bright field area, the R7 population was easily distinguishable from the other subpopulations in both culture conditions (Supplementary Fig. 8), further confirming the R7 subpopulation as mature MKs.

### Enrichment and isolation of a repopulating MKRP progenitor cell population

The ability to identify and isolate a repopulating MKRP cell population in an easy and high throughput fashion would significantly improve studies on MK and platelet development. To investigate whether LSK surface expression markers can be used to predict MK cell development, PCA was recalculated using the full set of morphological features and the principal component projections of the no TPO LSK experimental group were re-colored according to the expression of c-Kit and Sca-1 (Fig. 6A, Supplementary Methods). Using PCA, the R4 subpopulation of cells can be distinguished from the other subpopulations, which have greater c-Kit expression and extend further along the PC1 axis, while the Sca-1 expression does not extend as uniformly along the PC2 axis (Fig. 6B,C). Because the area feature negatively correlates with PC1 (Fig. 6B, Supplementary Fig. 4D), R4 cells with high c-Kit expression and corresponding low PC1 scores, are more likely to be larger in size compared to the other subpopulations (Supplementary Fig. 4). Similarly, because the feature circularity is negatively correlated with PC2, the R4 subpopulation of cells with low Sca-1 expression and corresponding low PC2 scores are more likely to be circular in shape (Supplementary Fig. 4D). Performing the same analysis for the cells grown in TPO amplifies the trends observed in the no TPO experimental group and results in a stronger separation between the four subpopulations (Fig. 6C, Supplementary Fig. 4E). Taken together, these results indicate that the R4 subpopulation of cells with similar c-Kit/Sca-1 expression also have similar morphologies, and that culturing with TPO exaggerates both the expression of these markers and the morphologies to which they correlate.

**Figure 6:**
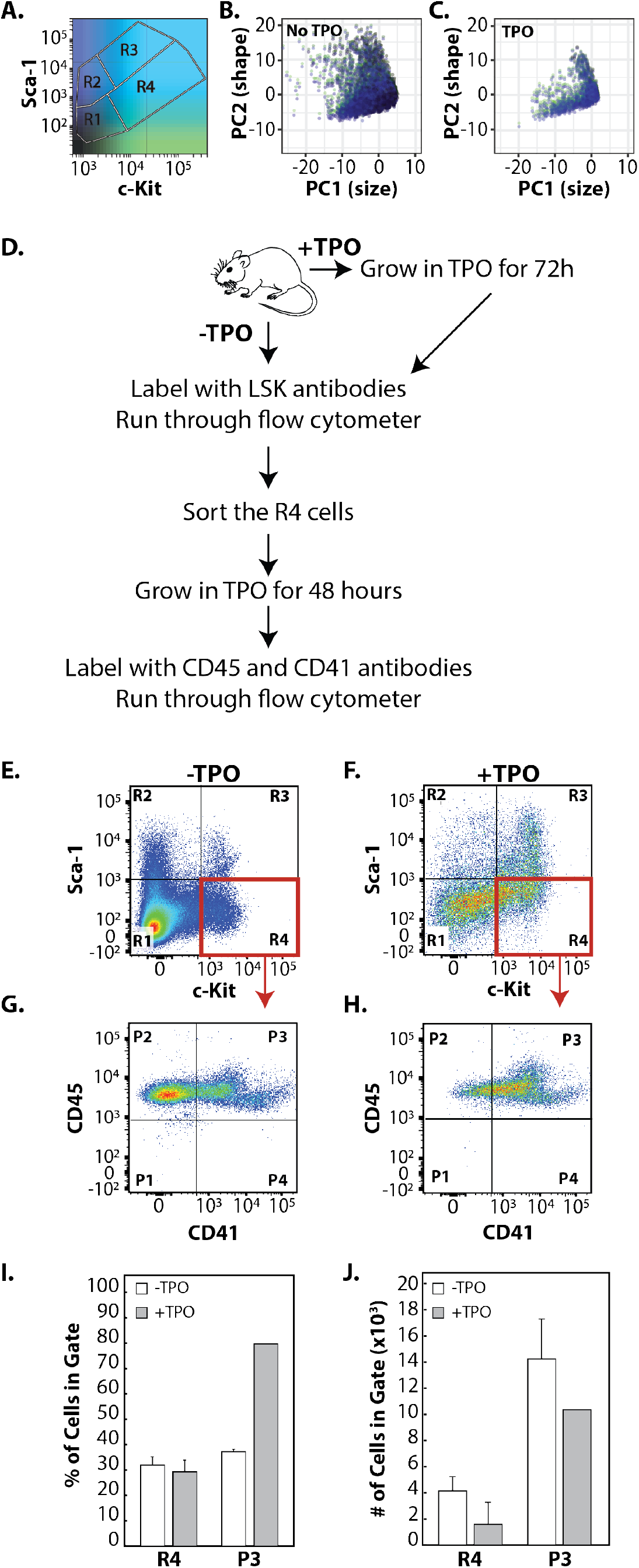
Identification of a repopulating MKRP cell population. **A.** Colormap with the R1-R4 subpopulation mask superimposed on top of cells directly harvested from the bone marrow (no TPO), stained for LSK antibodies, and run on the flow cytometer. **B.** PCA 1 and 2 projections of no TPO cells and **C.** of TPO cells using full sets of morphological features. **D.** Overview of experimental design. Bone marrow cells were harvested and split into two populations: 1. Cells initially not exposed to TPO were labeled with LSK antibodies, run on the flow cytometer, and the R4 subpopulation was sorted and grown in TPO before labeling with the MK cell markers, CD45 and CD41. 2. Cells initially grown in TPO for 72 hours, then labeled with LSK antibodies, run on the flow cytometer, and the R4 subpopulation was sorted and regrown in TPO for 48 hours, and stained with the MK cell markers CD45 and CD41. **E.** Flow cytometry plot of cells isolated directly from the bone marrow, stained for LSK antibodies and run on the flow cytometer. The defined subpopulations were identified (R1-R4). **F.** Flow cytometry plot of cells after being removed from the bone marrow, grown in TPO for 72 hours, stained for LSK antibodies, run on the flow cytometer and the defined subpopulations were identified (R1-R4). **G-H.** Cells from the R4 subpopulations (red box in E,F) of each culture condition were sorted, grown in TPO for 48 hours, and stained for the MK membrane markers CD45 and CD41 and run on the flow cytometer **G.** without priming cells with TPO, and **H.** with priming cells with TPO. New subpopulations were identified (P1-P4) by the expression of MK surface markers. **I.** Percentage of cells in the R4 (MKRP cells) and P4 (MKs) not primed **J.** Number of cells in the R4 (MKRP cells) and P4 (MKs) primed with TPO. For **I** and **J**, error bars represent the standard deviation of the mean from three independent bone marrow harvests from mice.

PCA analysis strongly suggests that the R4 subpopulation of LSK cells potentially contain a repopulating MK progenitor cell population. To directly test this, the −TPO population of cells were isolated from the bone marrow, washed, and immediately stained with LSK antibodies. The defined R4 subpopulation of cells were sorted using fluorescent activated cell sorting (FACS), then grown with TPO for 48 hours, labeled with MK antibodies, and the expression of these cell surface markers were assessed on the flow cytometer. In the +TPO group, cells were harvested from the bone marrow, washed and grown in the presence of TPO for 72 hours. Next, these cells were stained with LSK antibodies and the defined R4 subpopulation of cells were sorted using FACS, then grown with TPO for an additional 48 hours. The +TPO cells were then labeled with MK antibodies and the expression of these cell surface markers were assessed on the flow cytometer (Fig. 6D). In the −TPO group, it was found that 30% of cells were c-Kit^+^ (Fig. 6E,I), consistent with our previous results. When the R4 cells from the −TPO group were sorted using FACS, then cultured with TPO for an additional 48 hours, it was found that the cells continued to proliferate, but only 30% of the cells differentiated in MKs (Fig. 6G,I,J). Also consistent with previous results, 30% of cells in the +TPO group remained c-Kit^+^ after the first 72 hours of culture with TPO (Fig. 6F). However, 80% of the R4 cells from the +TPO group differentiated into MKs (Fig. 6H,I) while continuing to proliferate (Fig. 6J). These data demonstrate that cells pre-treated with TPO had a much higher propensity to differentiate into mature MKs compared to the cells not pre-treated with TPO, and indicates TPO pre-treatment as a method for the identification and enrichment of an MKRP cell population capable of self-renewal.

## Discussion

The classic model of hematopoiesis has been challenged over the last several years, transitioning away from the traditional, sequential differentiation pathway toward one that includes branches, bypassing and feedback loops^12,13,16,21,29^. One of the most highly debated aspects of the continuously evolving hematopoietic cell lineage organization is the origin and development of unipotent MKRP cells capable of giving rise to mature MKs. LT-HSCs and MKs have many features in common; most appreciably are their mutual dependence on TPO/c-Mpl stimulation and surface marker expression^5,8,16,19–22,30^. As a result, drawing the line between LT-HSCs, MKRP cells, and mature MKs to systematically study these cells has been exceedingly difficult. Previous methods have lacked the ability to image cells while simultaneously assessing the surface marker expression in response to culture conditions, ultimately limiting analytical capabilities. This study coupled cell surface antibody labeling and image flow cytometry to collect morphology data that is intrinsically coupled with surface marker expression to identify elusive MKRP cell types that can be further isolated to study the mechanisms of MK maturation and platelet development.

By combining mathematical approaches with experimental design, we identified, enriched, and isolated a population of MKRP cells. Subsequent characterization of these cells revealed that the c-Kit^+^Sca-1^−^Lin^−^ fraction of cultured bone marrow cells was highly enriched for unipotent LT-HSC-derived MKRP cells as defined by the CD9^+^ CD41^+^ c-Kit^+^ Sca-1^−^ IL7Ra^−^ Thy1.1^−^Lin^−^ immunotype. Furthermore, sorted MKRP cells that had been previously exposed to TPO retained the ability to self-renew and readily differentiate into mature MKs when re-cultured in TPO. Because these cells continue to proliferate when recultured with TPO after FACS, they can be considered a LT-HSC that has gone through the myeloid bypass pathway of differentiation to become MKRP cells.

Altogether, this study validates the use of minimal antibody panels and simple cell culture techniques for the enrichment and isolation of MKRP cells. Applying these methods to the disease setting may improve insights to the early stages of MK commitment and development. Additionally, these methods can be used to better understand the molecular mechanisms underlying atypical MK and platelet development to improve targeted therapies for diseases that arise from skewed cell fate decisions. Finally, this work sets the stage to better understand the role of dynamic gene expression patterns during the development of MKs and platelets to inform alternative approaches for directing stem cell fate that may include reprogramming progenitor cells with gene circuits to drastically increase the production of MKs and platelets *in vitro* ^31–34^.

## Acknowledgements

This work was funded by the National Science Foundation CAREER Program (CBET-1554017), the Office of Naval Research Young Investigator Program (N00014-16-1-3012), the National Institutes of Health Trailblazer Award (1R21EB025413-01), and the National Institutes of Health Director’s New Innovator Award (1DP2CA250006-01). T.L.D. would also like to acknowledge support by the University of Utah Flow Cytometry Facility in addition to the National Cancer Institute through Award Number 5P30CA042014-24. Research reported in this publication was supported by the National Center for Research Resources of the National Institutes of Health under Award Number 1S10RR026802-01. We also thank Dr. Jeff Weiss (U. Utah) for helpful discussions about PCA.

## Authors’ Contributions

LMB conducted all of the experiments. CHP performed the principal components analysis. LMB, JEM, and TLD designed the experiments and analyzed the data. LMB, CHP, and TLD wrote the manuscript.

## Data Availability

Data in used in the PCA in this manuscript were obtained from the ImageStream^®^X Mark II Imaging flow cytometer, as detailed in the Supplementary Methods. We will freely share data and code upon request.

## SUPPLEMENTARY INFORMATION

### Supplementary Figures

**Supplementary Figure 1.**
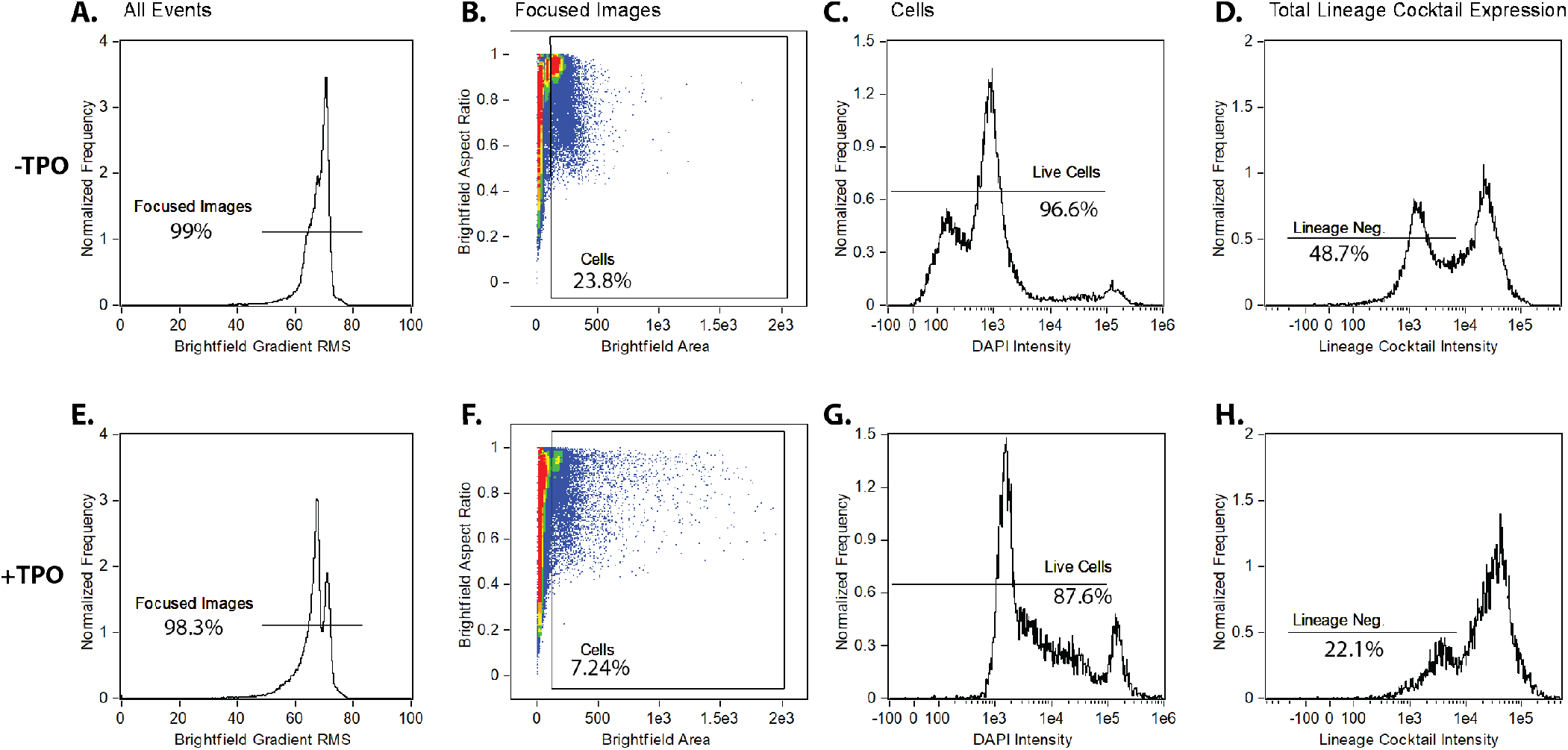
Gating strategy for Figure 2. **A.** Histogram of all events recorded by the ImageStream^®^X Mark II Imaging flow cytometer, gated for focused images. **B.** Dot plot of focused images gated for cells. **C.** Histogram of cells gated on DAPI intensity to identify live cells. **D.** Histogram of live cells gated on lineage cocktail intensity with a lineage negative gate indicating the cells not stained with lineage cocktail markers. Top row: cells are stained immediately after isolation from the mouse bone marrow. Bottom row: cells cultured with 50 ng/mL TPO for three days.

**Supplementary Figure 2.**
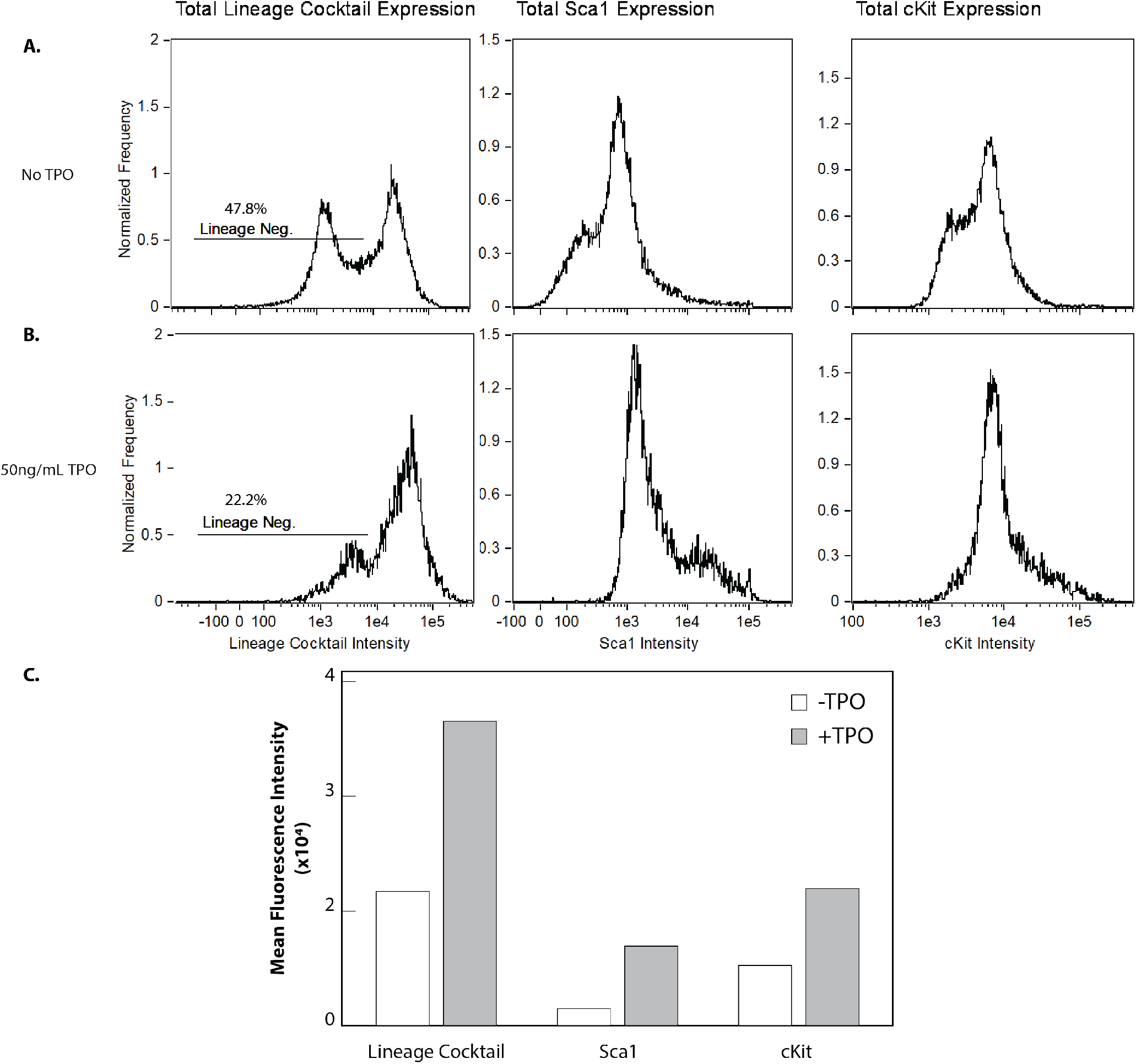
Total surface marker expression of bone marrow cells stained with lineage cocktail, Sca-1, and c-Kit before and after TPO exposure. **A.** Mean fluorescence intensities (MFI) of lineage cocktail, Sca-1, and c-Kit antibody staining of bone marrow cells directly isolated from the mouse. **B.** MFI of lineage cocktail, Sca-1 and c-Kit antibody staining of bone marrow cells cultured with 50 ng/mL TPO for 72 hours. **C.** Bar graph summarizing the data in **A-B**.

**Supplementary Figure 3.**
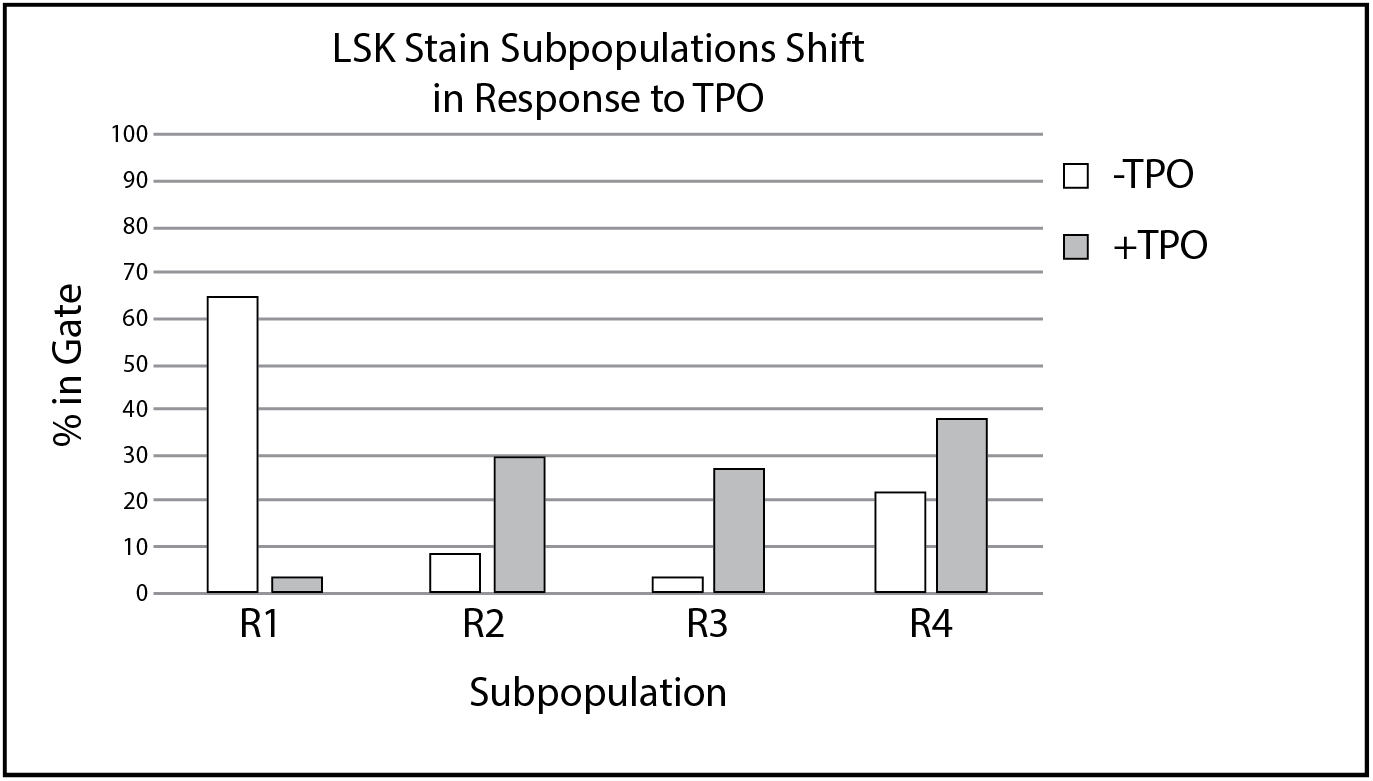
Quantification of lineage negative cells in each subpopulation. Percent of gated cells in R1-R4 subpopulations in Figure 1. The percentage of cells in R1 decreases, while the percentage of cells in R2-R4 increases, respectively, in response to culturing with TPO.

**Supplementary Figure 4.**
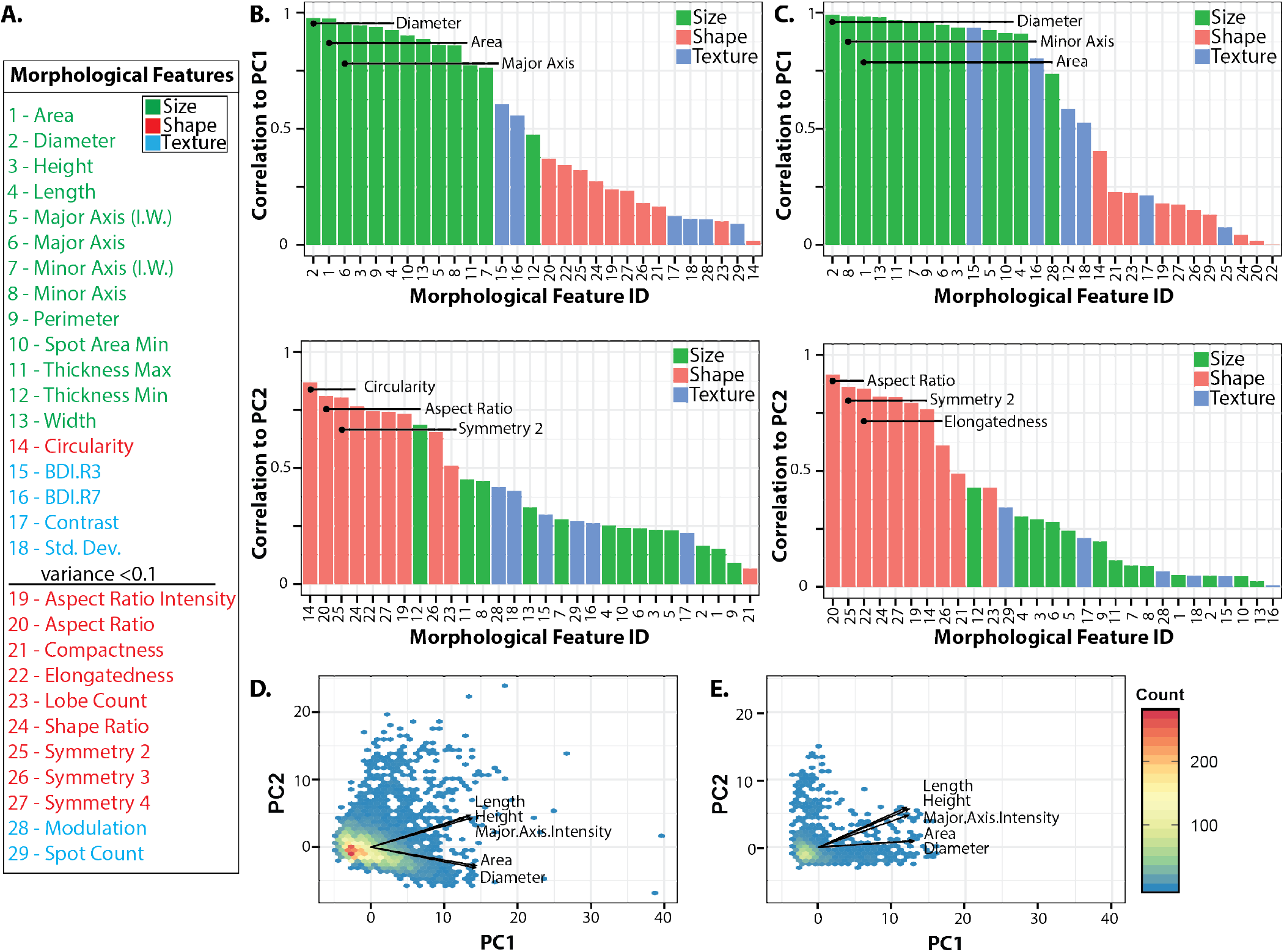
Supplementary PCA Data. **A**. Morphological feature list ID. Color indicates feature type: size (green), shape (red), and texture (blue). **B.** The absolute contribution/correlation of each morphological feature to PC1 and PC2 without TPO **C.** and with TPO using the full set of morphological features. The top 3 most highly correlated features are listed for each principal component. **D.** Principal component projections of LSK grown without TPO and **E.** with TPO. **D-E** were recolored based on their relative c-Kit (green) and Sca-1 (blue) expression values.

**Supplementary Figure 5.**
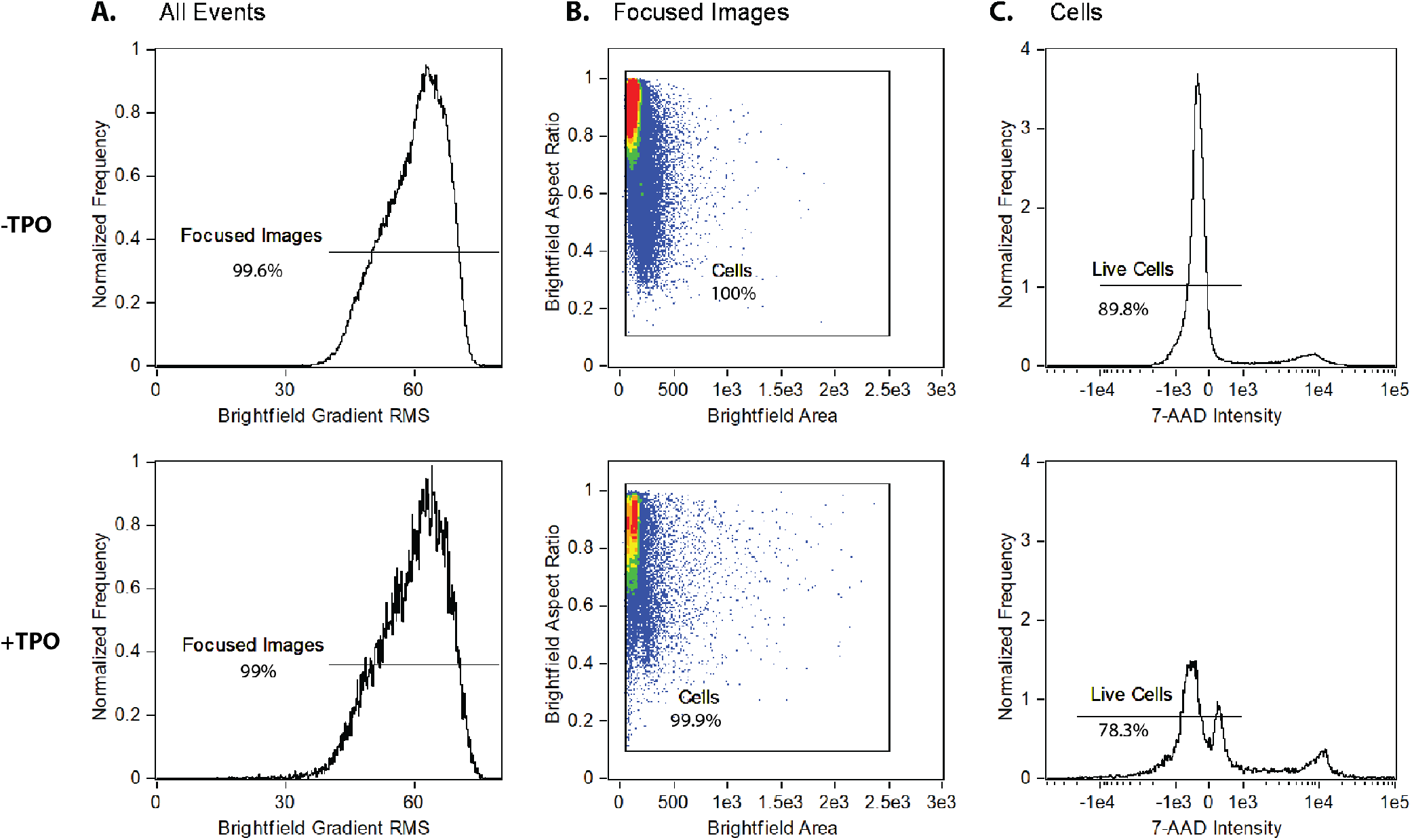
Gating strategy for Figure 5. **A.** Histogram of all events recorded by the ImageStream^®^X Mark II Imaging flow cytometer, gated for focused images. **B.** Dot plot of focused images gated for cells. **C.** Histogram of cells gated on 7-AAD intensity to identify live cells. Top row: cells are stained immediately after isolation from the mouse bone marrow. Bottom row: cells cultured with 50 ng/mL TPO for three days.

**Supplementary Figure 6.**
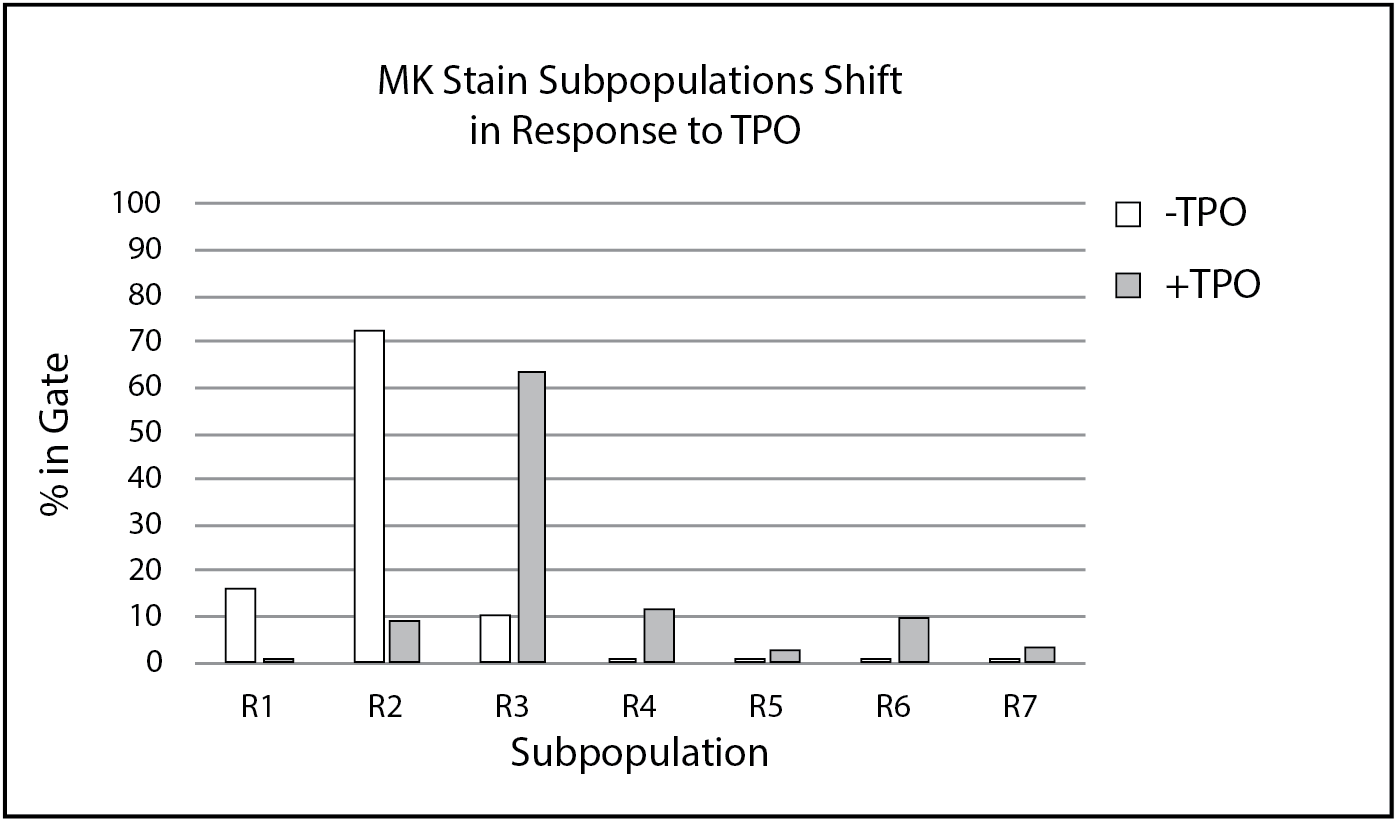
Quantification of live cells in each subpopulation. Percent of gated cells in R1-R7 subpopulations from Figure 5. The percentage of cells in R1-R2 decreases, while the percentage of cells in R2-R7 increases, respectively, in response to culturing with TPO.

**Supplementary Figure 7.**
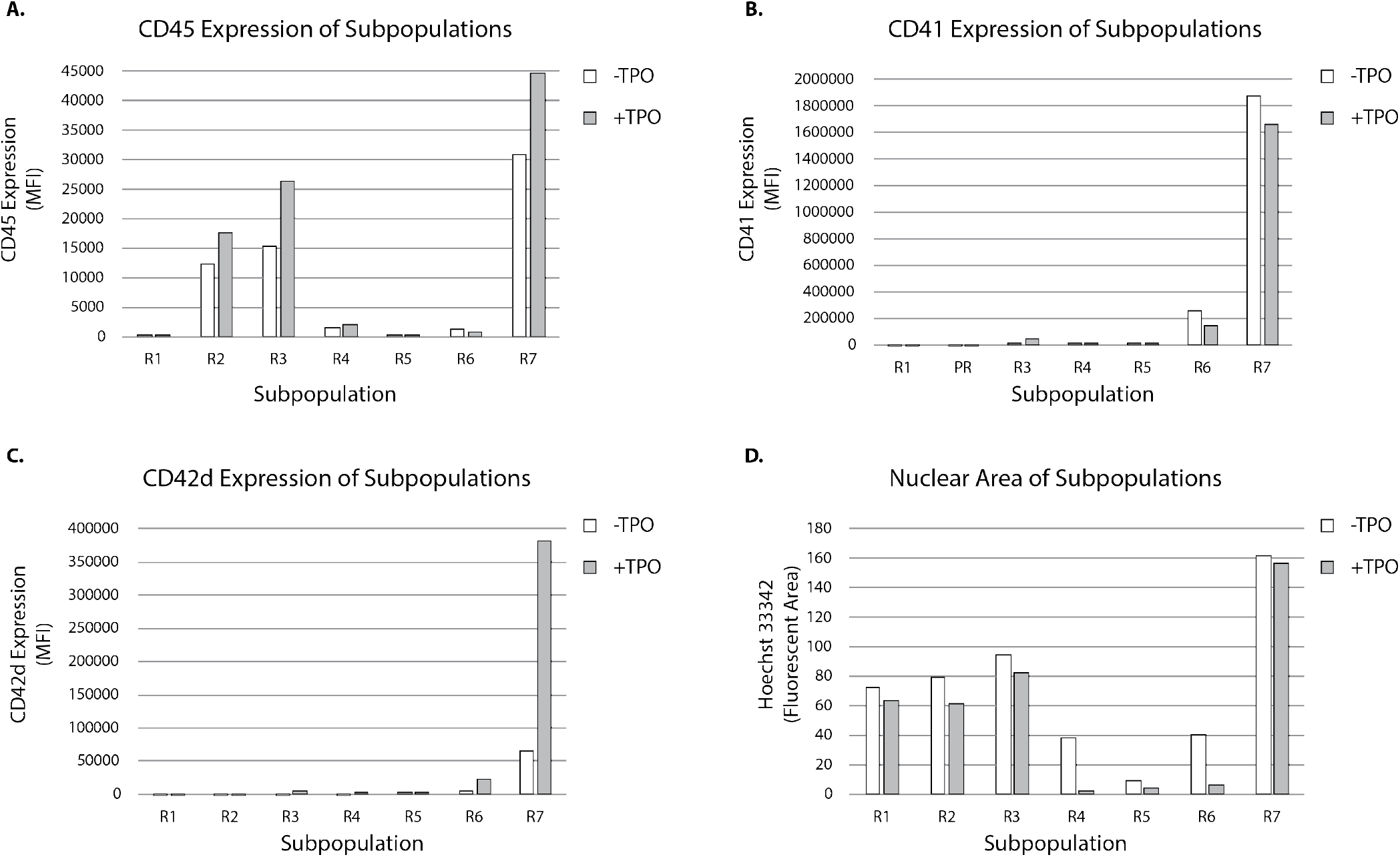
Analysis of surface marker expression and DNA content of subpopulations in Figure 5. **A.** MFI of CD45 positive cells in each subpopulation before and after TPO exposure. **B.** MFI of CD41 positive cells in each subpopulation before and after TPO exposure. **C.** MFI of CD42d positive cells in each subpopulation before and after TPO exposure. **D.** Nuclear area of cells in each population before and after TPO exposure.

**Supplementary Figure 8.**
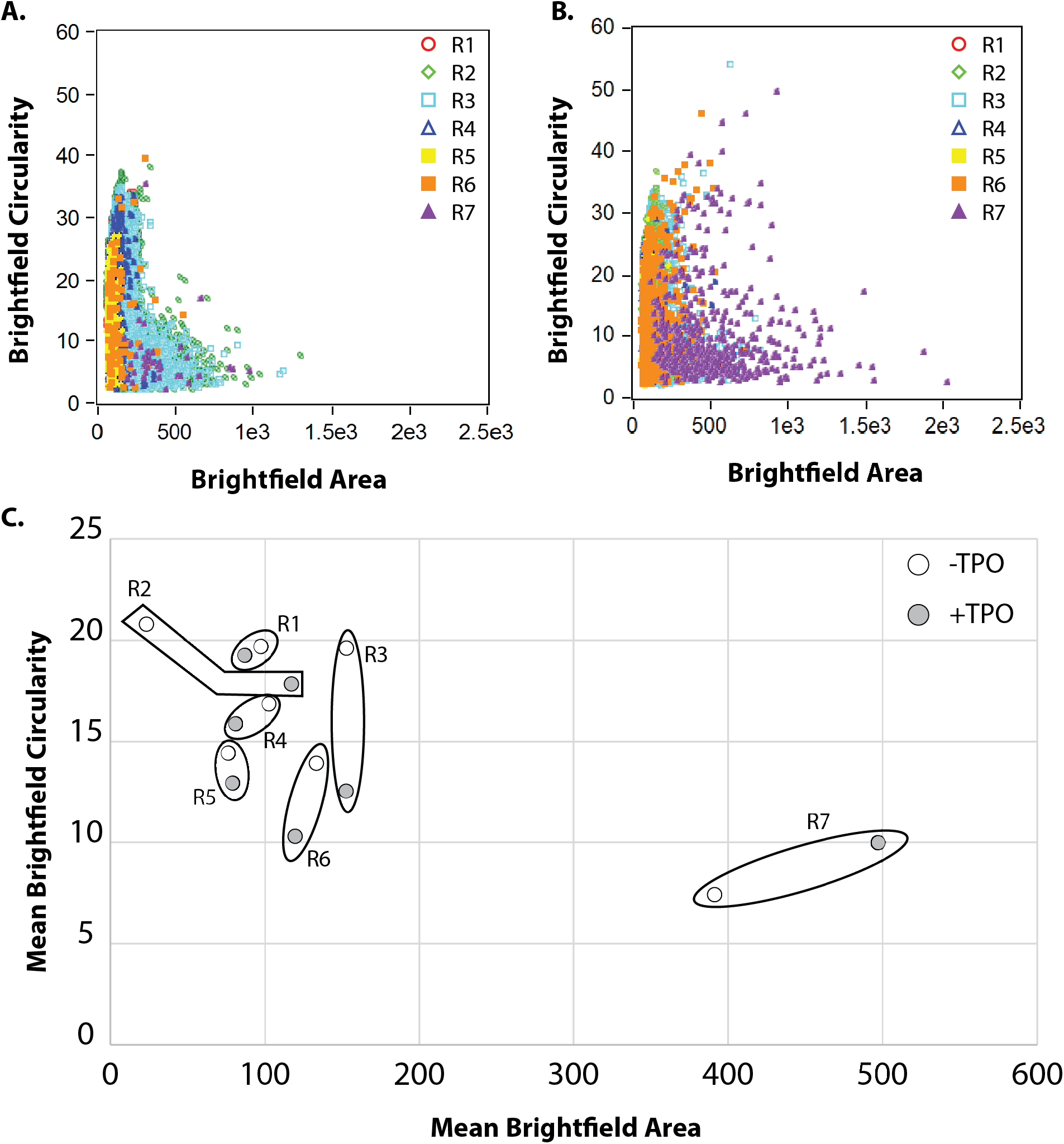
Brightfield circularity versus areas plots for each subpopulation enables identification of MKs by cell size alone. **A.** Brightfield circularity vs. bright field area plot of cells directly isolated from the mouse bone marrow and not exposed to TPO. **B.** Brightfield circularity vs. bright field area plots of cells after culture with 50 ng/mL of TPO. **C.** Mean brightfield circularity vs. brightfield area of the different subpopulations for the two culture conditions.

### Supplementary Table

**Table S1:** Reagents used in flow cytometry experiments. A comprehensive list of reagents used in each of the flow cytometry experiments, including reagent name, specificity/target information, brand, clone (if applicable), catalog and/or reference number, and dilution.

| Key Resources Table |  |  |  |  |
| --- | --- | --- | --- | --- |
| Reagent Type (species) or resource | Designation | Source or Reference | Identifiers | Additional Information |
| Antibody | PE-Cy<sup>TM</sup>7 Rat anti-Mouse CD117 | BD Pharmingen | 561681 | Clone 2B8; Dilution used = 1:133 |
| Antibody | PE Rat Anti-Mouse Ly-6A/E | BD Pharmingen | 562059 | Clone D7; Dilution used = 1:100 |
| Antibody | APC Mouse Lineage Antibody Cocktail | BD Pharmingen | 51-9003632 | Dilution used = 1:5 |
| Antibody | CD45 Monoclonal Antibody, PE-Cyanine7, eBioscience<sup>TM</sup> | eBioscience | 25-0451-81 | Clone 30-F11; Dilution used = 1:133 |
| Antibody | PE Rat Anti-Mouse CD41 | BD Pharmingen | 561850 | Clone MWReg30; Dilution used = 1:100 |
| Antibody | CD42d Monoclonal Antibody, APC, eBioscience<sup>TM</sup> | eBioscience | 17-0421-80 | Clone 1C2; Dilution used = 1:50 |
| Antibody | BV421 Rat Anti-Mouse CD117 | BD Biosciences | 566290 | Clone 2B8; Dilution used = 1:80 |
| Antibody | BV510 Rat Anti-Mouse CD9 | BD Biosciences | 740128 | Clone KMC8; Dilution used = 1:80 |
| Antibody | CD90.1 (Thy-1.1) Monoclonal Antibody (HIS51), PerCP-Cyanine5.5, eBioscience<sup>TM</sup> | ThermoFisher Scientific | 45-0900-80 | Clone HIS51; Dilution used = 1:333 |
| Antibody | CD41a Monoclonal Antibody (eBioMWReg30 (MWReg30)), PE-Cyanine7, eBioscience<sup>TM</sup> | ThermoFisher Scientific | 25-0411-80 | Clone MWReg30; Dilution used = 1:60 |
| Antibody | CD127 Monoclonal Antibody (A7R34), APC-eFluor 780, eBioscience<sup>TM</sup> | ThermoFisher Scientific | 47-1271-80 | Clone A7R34; Dilution used = 1:80 |
| DNA Dye | DAPI | Thermo Scientific | 62247 |  |
| DNA Dye | Hoechst 33342 Solution | BD Pharmingen | 561908 | Dilution used = 1:1000 |
| DNA Dye | eBioscience<sup>TM</sup> 7-AAD Viability Staining Solution | eBioscience | 00-6993-50 |  |
| DNA dye | SYTOX™ Green Dead Cell Stain, for flow cytometry | ThermoFisher Scientific | S34860 |  |
| Equipment | ImageStream®X Mark II Imaging Flow Cytometer | Merck Millipore | NA |  |
| Equipment | BD FACSAria™ III | BD Biosciences | NA |  |
| Equipment | BD FACSCanto™ II | BD Biosciences | NA |  |
| Other | Fetal Bovine Serum | Gibco | 16000044 |  |
| Other | Penicillin Streptomycin | ThermoFisher Scientific | 15140122 |  |
| Other | IMDM Medium | ThermoFisher Scientific | 12440053 |  |
| Other | 26 G BD™ Needle 1/2 in. single use, sterile | BD | 305111 |  |
| Other | 35mm Corning™ TC-Treated Culture Dishes | Fisher Scientific | 08-772-20 |  |
| Other | ACK Lysis Buffer | ThermoFisher Scientific | A1049201 |  |
| Other | HBSS | Gibco | 14025092 |  |
| Other | Bovine Serum Albumin | ThermoFisher Scientific | BP1600-100 |  |
| Peptide, recombinant protein | Recombinant murine TPO | Peprtech | 315-14 |  |
| Software | IDEAS® | Amnis Corporation | NA |  |
| Software | FlowJo® | FlowJo, LLC | NA |  |
| Strain, (*Mus musculus*) | C57BL/6 | NA | NA |  |

## SUPPLEMENTAL METHODS

### Materials and Methods

#### Bone marrow dissection and cell culture

Mice were euthanatized by CO2 asphyxiation, followed by cervical dislocation based upon IACUC approved guidelines. Femurs were dissected from the mouse, excess tissue was removed, and the epiphyses were snipped off. Bone marrow was flushed with IMDM (ThermoFisher Scientific, #12440053) + 10% FBS (Gibco, #16000044) and 1X Penicillin-Streptomycin (ThermoFisher Scientific, #15140122) using a 26.5-gauge needle and collected in a 35mm dish. Red blood cells were lysed using ACK Lysis Buffer (ThermoFisher Scientific, # A1049201) for 5 minutes at room temperature, then the solution was centrifuged at 300x g for 5 minutes. The cells were washed 2x with HBSS (Gibco, #14025092) then counted using a hemocytometer. If cells were cultured before analysis, 1×10^7^ cells were incubated in IMDM + 10% FBS supplemented with 50 ng/mL of recombinant murine thrombopoietin (TPO) (PeproTech, #315-14) in a T75 flask at 37°C in 5% CO_2_ for 72 hours.

#### Preparation for image flow cytometry

After isolation from the bone marrow or culture with TPO for 72 h, cells were washed 2x with PBS. For the LSK stain, cells were resuspended in PBS + 3% BSA (ThermoFisher Scientific, #BP1600-100) with the addition of PE-Cy™7 Rat anti-Mouse CD117, PE Rat Anti-Mouse Ly-6A/E, and APC Mouse Lineage Antibody Cocktail, and incubated on ice protected from light for 30 min. Cells were washed 2x with PBS and resuspended at a final concentration of 2×107 cells/ml in PBS. DAPI was added at least 5 min prior to running the sample on the image flow cytometer.

For the MK stain, cells were resuspended in prewarmed IMDM + 10% FBS containing Hoechst 33342 solution, then incubated at 37°C for 45 minutes. Cells were washed with PBS then resuspended in PBS + 3% BSA with the addition of PE-Cyanine7 CD45 Monoclonal Antibody, PE Rat Anti-Mouse CD41, and APC CD42d Monoclonal Antibody and incubated on ice, protected from light for 30 minutes. Cells were washed 2x with PBS then resuspended at a final concentration of 2×10^7^ cells/mL in PBS. 7-AAD was added at least 5 minutes prior to running the sample on the image flow cytometer. For a complete description of staining reagents, see Supplementary Table 1.

#### Image flow cytometry data acquisition and analysis

All experiments used the ImageStream®X Mark II Imaging Flow Cytometer and all image flow cytometry analysis was performed in the IDEAS® Software. Data from image flow cytometry experiments was acquired by gating for events that were in focus, did not contain calibration focus beads, and excluded viability dyes. For a full description of data acquisition and analysis methods, consult the INSPIRE™ ImageStream®X Mark II Imaging Flow Cytometer user’s manual and the IDEAS® software user manual available online. Gating strategies used in all experiments are described in the Supplemental Information (Supplementary Fig. 1,5).

#### Image Analysis and Morphological Feature Extraction

With the LSK stain dataset, brightfield images of lineage-negative cells from each experimental group (no TPO and treated with 50 ng/mL TPO) were first isolated from the full dataset. Individual cells in each image were distinguished from the background and defined automatically using the default brightfield settings in the IDEAS® software to produce masked images. Next, the IDEAS® software’s built-in feature analysis utility was applied to the masked images and used to measure various morphological features of each cell. In total, twenty-nine morphological features were computed for each cell. Thirteen size features were computed: area; diameter, height; length; major/minor axis; major/minor axis weighted by intensity; perimeter; spot area min; thickness min/max; and width. Next, ten shape features were computed: aspect ratio; intensity weighted aspect ratio; circularity; compactness; elongatedness; lobe count; shape ratio; two-lobe symmetry (symmetry 2); three lobe symmetry (symmetry 3); and four lobe symmetry (symmetry 4). Lastly, six texture features were computed: the intensity of bright details with radii less than 3 (BDI.R3) and radii less than 7 (BDI.R7); contrast; modulation; spot count; and standard deviation. Full descriptions of these features and the means by which they are computed can be found in the IDEAS® software user manual. Finally, each observed cell was assigned a categorical variable, R1, R2, R3, or R4, based on the previously specified gates for Sca-1 and c-Kit expression. For the surface marker expression analysis, each cell was assigned its corresponding Sca-1 and c-Kit expression level instead of a categorical variable.

#### Principal Components Analysis

The variance of all twenty-nine morphological features was computed and those with variances less than 0.1 were omitted from further analysis (Supplementary Fig. 4). These features were aspect ratio intensity, aspect ratio, compactness, elongatedness, lobe count, shape ratio, symmetry 2-4, modulation, and spot count. The remaining eighteen morphological features from 31,265 observations in the no TPO dataset were centered, and scaled/normalized such that the values of each feature ranged from 0 to 1 (i.e. min-max normalization). Next, the principal components of the normalized dataset were computed using R’s **prcomp** function^1^. The number of principal components needed to describe each dataset was determined using the broken stick model and David Zelený‘s **evplot** function^2,3^. Observed principal components that describe more variance than the amount of variance predicted under the broken-stick model are considered necessary to effectively interpret the data. In this experiment, it was determined that the first two principal components were needed to adequately describe the data. Finally, bi-plots (plots with principal components on each axes) and correlation plots were generated with the **ggplot2** library^4^. These steps were repeated for the 50 ng/mL TPO dataset to generate the biplot (Fig. 3 B, D, F). For the surface marker expression bi-plots, the color of each observation was determined based on the relative expression of the surface markers c-Kit (green) and Sca-1 (blue) (Fig. 4, Supplementary Fig. 4). The dynamic range of this color map was defined to capture 99% of the observations. These same steps were repeated to generate the principal components for the 10,926 observations from the 50 ng/mL dataset.

#### Fluorescence activated cell sorting (FACS), re-culture, and antibody staining of the R4 cell subpopulation

For the −TPO group, cells were directly isolated from the mouse bone marrow and prepared for FACS using LSK antibodies as described above. For the +TPO group, cells were isolated from the mouse bone marrow and cultured with 50 ng/mL of TPO for 3 days, then prepared for FACS using LSK antibodies as described above. Cells were then sorted into IMDM + 10% FBS with the addition of 50 ng/mL TPO using the BD FACSAria™ III, then plated into a cell culture dish and incubated at 37°C for an additional 48 hours. After 48 hours, cells were stained with PE-Cyanine7 CD45 monoclonal antibody and PE rat anti-mouse CD41 and run on the BD FACSCanto™ II. All BD FACSAria™ III and BD FACSCanto™ II data analysis was performed using FlowJo®.

## References

1 Laffont, B. et al. Activated platelets can deliver mRNA regulatory Ago2*microRNA complexes to endothelial cells via microparticles. Blood 122, 253–261, doi:10.1182/blood-2013-03-492801 (2013).

2 Leiter, O. & Walker, T. L. Platelets: The missing link between the blood and brain? Prog Neurobiol 183, 101695, doi:10.1016/j.pneurobio.2019.101695 (2019).

3 Nishikii, H., Kurita, N. & Chiba, S. The Road Map for Megakaryopoietic Lineage from Hematopoietic Stem/Progenitor Cells. Stem Cells Transl Med 6, 1661–1665, doi:10.1002/sctm.16-0490 (2017).

4 Kuter, D. J., Beeler, D. L. & Rosenberg, R. D. The purification of megapoietin: a physiological regulator of megakaryocyte growth and platelet production. Proceedings of the National Academy of Sciences of the United States of America 91, 11104–11108 (1994).

5 Machlus, K. R. & Italiano, J. E., Jr. The incredible journey: From megakaryocyte development to platelet formation. The Journal of cell biology 201, 785–796, doi:10.1083/jcb.201304054 (2013).

6 Patel, S. R., Hartwig, J. H. & Italiano, J. E., Jr. The biogenesis of platelets from megakaryocyte proplatelets. The Journal of clinical investigation 115, 3348–3354, doi:10.1172/JCI26891 (2005).

7 Thon, J. N. & Italiano, J. E. Platelet formation. Seminars in hematology 47, 220–226, doi:10.1053/j.seminhematol.2010.03.005 (2010).

8 Kaushansky, K. et al. Promotion of megakaryocyte progenitor expansion and differentiation by the c-Mpl ligand thrombopoietin. Nature 369, 568–571, doi:10.1038/369568a0 (1994).

9 Xavier-Ferrucio, J. & Krause, D. S. Concise Review: Bipotent Megakaryocytic-Erythroid Progenitors: Concepts and Controversies. Stem Cells 36, 1138–1145, doi:10.1002/stem.2834 (2018).

10 Nakeff, A. & Maat, B. Separation of megakaryocytes from mouse bone marrow by velocity sedimentation. Blood 43, 591–595 (1974).

11 Wilson, A. & Trumpp, A. Bone-marrow haematopoietic-stem-cell niches. Nature reviews. Immunology 6, 93–106, doi:10.1038/nri1779 (2006).

12 Yamamoto, R. et al. Clonal analysis unveils self-renewing lineage-restricted progenitors generated directly from hematopoietic stem cells. Cell 154, 1112–1126, doi:10.1016/j.cell.2013.08.007 (2013).

13 Paul, F. et al. Transcriptional Heterogeneity and Lineage Commitment in Myeloid Progenitors. Cell 163, 1663–1677, doi:10.1016/j.cell.2015.11.013 (2015).

14 Woolthuis, C. M. & Park, C. Y. Hematopoietic stem/progenitor cell commitment to the megakaryocyte lineage. Blood 127, 1242–1248, doi:10.1182/blood-2015-07-607945 (2016).

15 Noetzli, L. J., French, S. L. & Machlus, K. R. New Insights Into the Differentiation of Megakaryocytes From Hematopoietic Progenitors. Arteriosclerosis, thrombosis, and vascular biology 39, 1288–1300, doi:10.1161/ATVBAHA.119.312129 (2019).

16 Chang, Y., Bluteau, D., Debili, N. & Vainchenker, W. From hematopoietic stem cells to platelets. Journal of thrombosis and haemostasis : JTH 5 Suppl 1, 318–327, doi:10.1111/j.1538-7836.2007.02472.x (2007).

17 Seita, J. & Weissman, I. L. Hematopoietic stem cell: self-renewal versus differentiation. Wiley Interdiscip Rev Syst Biol Med 2, 640–653, doi:10.1002/wsbm.86 (2010).

18 Buckman, C. et al. High throughput, parallel imaging and biomarker quantification of human spermatozoa by ImageStream flow cytometry. Syst Biol Reprod Med 55, 244–251, doi:10.3109/19396360903056224 (2009).

19 Buza-Vidas, N. et al. Cytokines regulate postnatal hematopoietic stem cell expansion: opposing roles of thrombopoietin and LNK. Genes & development 20, 2018–2023, doi:10.1101/gad.385606 (2006).

20 Kirito, K., Fox, N. & Kaushansky, K. Thrombopoietin stimulates Hoxb4 expression: an explanation for the favorable effects of TPO on hematopoietic stem cells. Blood 102, 3172–3178, doi:10.1182/blood-2003-03-0944 (2003).

21 Seita, J. et al. Lnk negatively regulates self-renewal of hematopoietic stem cells by modifying thrombopoietin-mediated signal transduction. Proceedings of the National Academy of Sciences of the United States of America 104, 2349–2354, doi:10.1073/pnas.0606238104 (2007).

22 Yoshihara, H. et al. Thrombopoietin/MPL signaling regulates hematopoietic stem cell quiescence and interaction with the osteoblastic niche. Cell Stem Cell 1, 685–697, doi:10.1016/j.stem.2007.10.020 (2007).

23 Ng, A. P. et al. Characterization of thrombopoietin (TPO)-responsive progenitor cells in adult mouse bone marrow with in vivo megakaryocyte and erythroid potential. Proceedings of the National Academy of Sciences of the United States of America 109, 2364–2369, doi:10.1073/pnas.1121385109 (2012).

24 Attema, J. L. et al. Epigenetic characterization of hematopoietic stem cell differentiation using miniChIP and bisulfite sequencing analysis. Proceedings of the National Academy of Sciences of the United States of America 104, 12371–12376, doi:10.1073/pnas.0704468104 (2007).

25 Wang, J. F., Liu, Z. Y. & Groopman, J. E. The alpha-chemokine receptor CXCR4 is expressed on the megakaryocytic lineage from progenitor to platelets and modulates migration and adhesion. Blood 92, 756–764 (1998).

26 Shin, J. Y., Hu, W., Naramura, M. & Park, C. Y. High c-Kit expression identifies hematopoietic stem cells with impaired self-renewal and megakaryocytic bias. The Journal of experimental medicine 211, 217–231, doi:10.1084/jem.20131128 (2014).

27 Nakorn, T. N., Miyamoto, T. & Weissman, I. L. Characterization of mouse clonogenic megakaryocyte progenitors. Proceedings of the National Academy of Sciences of the United States of America 100, 205–210, doi:10.1073/pnas.262655099 (2003).

28 Hashimoto, K. et al. Distinct hemogenic potential of endothelial cells and CD41+ cells in mouse embryos. Dev Growth Differ 49, 287–300, doi:10.1111/j.1440-169X.2007.00925.x (2007).

29 Reya, T., Morrison, S. J., Clarke, M. F. & Weissman, I. L. Stem cells, cancer, and cancer stem cells. Nature 414, 105–111, doi:10.1038/35102167 (2001).

30 Wickham, H. ggplot2: Elegant Graphics for Data Analysis. (Springer-Verlag New York, 2016).

31 Deans, T. L., Cantor, C. R. & Collins, J. J. A tunable genetic switch based on RNAi and repressor proteins for regulating gene expression in mammalian cells. Cell 130, 363–372, doi:https://doi.org/10.1016/j.cell.2007.05.045 (2007).

32 Fitzgerald, M., Gibbs, C., Shimpi, A. A. & Deans, T. L. Adoption of the Q Transcriptional System for Regulating Gene Expression in Stem Cells. ACS synthetic biology 6, 2014–2020, doi:https://doi.org/10.1021/acssynbio.7b00149 (2017).

33 Healy, C. P. & Deans, T. L. Genetic circuits to engineer tissues with alternative functions. J Biol Eng 13, 39, doi:10.1186/s13036-019-0170-7 (2019).

34 MacDonald, I. C. & Deans, T. L. Tools and applications in synthetic biology. Adv Drug Deliv Rev 105, 20–34, doi:10.1016/j.addr.2016.08.008 (2016).

## Supplementary References

1 Team, R. C. R: A Language and Environment for Statistical Computing <http://www.R-project.org/> (2018).

2 Zelený, D. Analysis of community ecology data in R, <https://www.davidzeleny.net/anadat-r/doku.php/en:numecolr:evplot> (2017).

3 Jackson, D. A. Stopping Rules in Principal Components Analysis: A Comparison of Heuristical and Statistical Approaches.Ecology 74, 2204–2214 (1993).

4 Wickham, H. ggplot2: Elegant Graphics for Data Analysis. (Springer-Verlag New York, 2016).

